# ViFIT-assisted Histopathology: From H&E Style Standardization to Virtual Fiber Image Transformation

**DOI:** 10.1101/2025.01.24.634654

**Authors:** Shu Wang, Xiao Zhang, Xingfu Wang, Chenyong Lv, Xiahui Han, Xiong Lin, Deyong Kang, Ruolan Lin, Liwen Hu, Feng Huang, Wenxi Liu, Jianxin Chen

## Abstract

Deep learning-based virtual fiber staining provides a promising complement to routine H&E pathology. However, the reliance on predefined staining style inputs and manual intervention limits the clinical applicability of existing methods. To address these challenges, we introduce ViFIT-assisted histopathology, a two-stage diagnostic approach that integrates our proposed unsupervised deep learning-based virtual fiber transformation model (ViFIT). This approach enables the conversion of H&E-stained images with diverse styles into pathologist-preferred H&E images, while simultaneously generating content-consistent virtual fiber images containing label-free collagen fibers and stained reticular and elastic fibers. ViFIT-assisted histopathology reveals tumor-associated fibers and provides quantitative metrics across multiple intraoperative and postoperative cases. Experimental results demonstrate that ViFIT significantly outperforms state-of-the-art unsupervised methods in both style standardization and virtual staining, across various downstream tasks and cancer types. By eliminating the need for staining variation and manual annotation, ViFIT-assisted histopathology streamlines histopathology workflows, making it well-suited for multi-center consultations and differential diagnosis.

## 1. Introduction

Histopathology is an essential component of gold-standard pathological diagnosis, playing an irreplaceable role in ensuring diagnostic accuracy and reliability^1^. Hematoxylin and eosin (H&E) staining is widely utilized in routine clinical pathology for the initial diagnosis, highlighting the morphological structures of nuclei and cytoplasm^2^. However, H&E staining fails to specifically capture certain extracellular matrix components, such as tumor-associated fibers, which undergo significant morphological changes during tumor progression. These alterations are critical for both differential diagnosis^3^ and prognostic prediction^4,5^. To clearly visualize these features, pathologists often resort to multiple specialized fiber staining for a comprehensive diagnosis^6–8^. Due to its stringent requirements for tissue fixation and staining protocol^9,10^, specialized staining techniques are challenging to implement in time-sensitive scenarios such as intraoperative diagnosis.

Deep learning-based virtual staining^11^, or virtual image transformation approaches, have recently attracted considerable attention from pathologists, as they offer a promising alternative by bypassing the labor-intensive and time-consuming staining process. These techniques leverage powerful deep generative neural networks to directly transform H&E-stained images into corresponding specifically stained images. However, their widespread application in pathology remains fundamentally limited by expert annotations and the scarcity of training data, particularly the lack of registered image pairs^12^, which hampers the image transformation capabilities. Consequently, unsupervised learning-based virtual staining approaches^13–15^ have emerged as a crucial area of research and constitute the focus of this study.

Despite its potential, unsupervised virtual staining confronts two significant challenges. Firstly, H&E images acquired from different institutions often exhibit diverse styles, which may result in variations in the representation of identical tissue features. Existing unsupervised transformation methods struggle to mitigate the negative impact of these style variations, thereby relying on input images with similar styles for effective virtual staining^16,17^. Notably, only supervised models have thus far attempted to tackle this issue. For instance, Su et al.^18^ employed Staintools for H&E image normalization prior to virtual immunohistochemical image transformation. However, these supervised models inherently require pixel-level registered datasets for training. Their heavy dependence on manual RGB color mapping for H&E images with minor stylistic differences restricts its generalization capabilities and entails significant labor costs.

Secondly, within the context of unsupervised learning, the model struggles to distinguish visually similar features during training without the guidance of supervisory signals. This often results in misclassification of similar cells or extracellular matrix components. Previous approaches often relied on manual interventions to alleviate this issue, such as requiring manual adjustments for color contrast enhancement^19^ or image content alignment^20^, which facilitated the discrimination of intricate tissue regions. Other studies first annotated features prone to misclassification and then trained an independent model to address these challenges. For instance, Liu et al.^21^ pre-trained a pathological representation network to identify cancerous lesion locations across domains. Likewise, Pati et al.^22^ introduced a cell classifier trained on expert pre-annotations of cellular staining status. However, these approaches remain heavily dependent on pathologists expertise, limiting their scalability.

To address the aforementioned limitations, we propose a novel deep learning-based unsupervised Virtual Fiber Image Transformation framework, denoted ViFIT, which is capable of transforming various styles of H&E-stained images into specific fiber images. ViFIT consists of two stages: H&E image style standardization and virtual fiber transformation. Specifically, ViFIT first employs additional pretext tasks to standardize diverse H&E staining styles while preserving image content. These standardized images, which retain significant diagnostic utility as intermediate outcomes, are further utilized to generate high-quality fiber representations through an intensity-reversed, fiber-sensitive module in a coarse-to-fine manner. Unlike existing approaches, our unsupervised framework requires no manual intervention in the two-stage process, thereby reducing the complexity associated with style variations and image pre-registration.

Furthermore, we introduce an efficient solution for auxiliary diagnostics, ViFIT-assisted histopathology, validated in both postoperative and intraoperative settings. It provides pathologists with preferred standardized H&E-stained images, generates various types of virtual fiber images, and reveals tumor-associated fiber patterns consistent with real-stained images. Furthermore, ViFIT-assisted histopathology offers quantitative metrics for collagen fiber alignment and orientation, reducing subjective variability and streamlining workflows. For evaluation, we establish a public H&E-to-fiber dataset that includes second-harmonic generation (SHG) collagen fibers, reticulin fiber-stained (RF), and Masson trichrome-stained (MT) images, spanning seven distinct tissue disease types. Comprehensive experimental results demonstrate that ViFIT outperforms the state-of-the-art methods for both unsupervised tasks of style standardization and virtual staining.

## 2. Results

### 2.1 ViFIT Workflow

In this work, we propose a novel framework, ViFIT, that aims to transform H&E images of diverse styles to virtual fiber images through two stages: H&E-stained image style standardization and virtual fiber transformation, as illustrated in Fig. 1. In the first stage, we design a generative adversarial network (GAN)-based model that can transform H&E images with various styles to the expert-consensus standard style without relying on any supervisory signals and image content tampering. The proposed model comprises of two cascaded generators, Standardization Generator (S-Gen) and Reconstruction Generator (R-Gen), both of which have the identical structure yet distinct parameters. During the training phase, like previous models^23^, a cyclic training scheme is applied. Specifically, given standard style images as references, the original H&E images are fed into S-Gen to generate standard style images, which are then subsequently delivered into R-Gen to reconstruct the original images. On the other hand, for cyclic training, the order of S-Gen and R-Gen is reversed. The standard style images are passed into R-Gen to produce H&E images with given styles, and then S-Gen attempts to standardize these images.

**Fig. 1.**
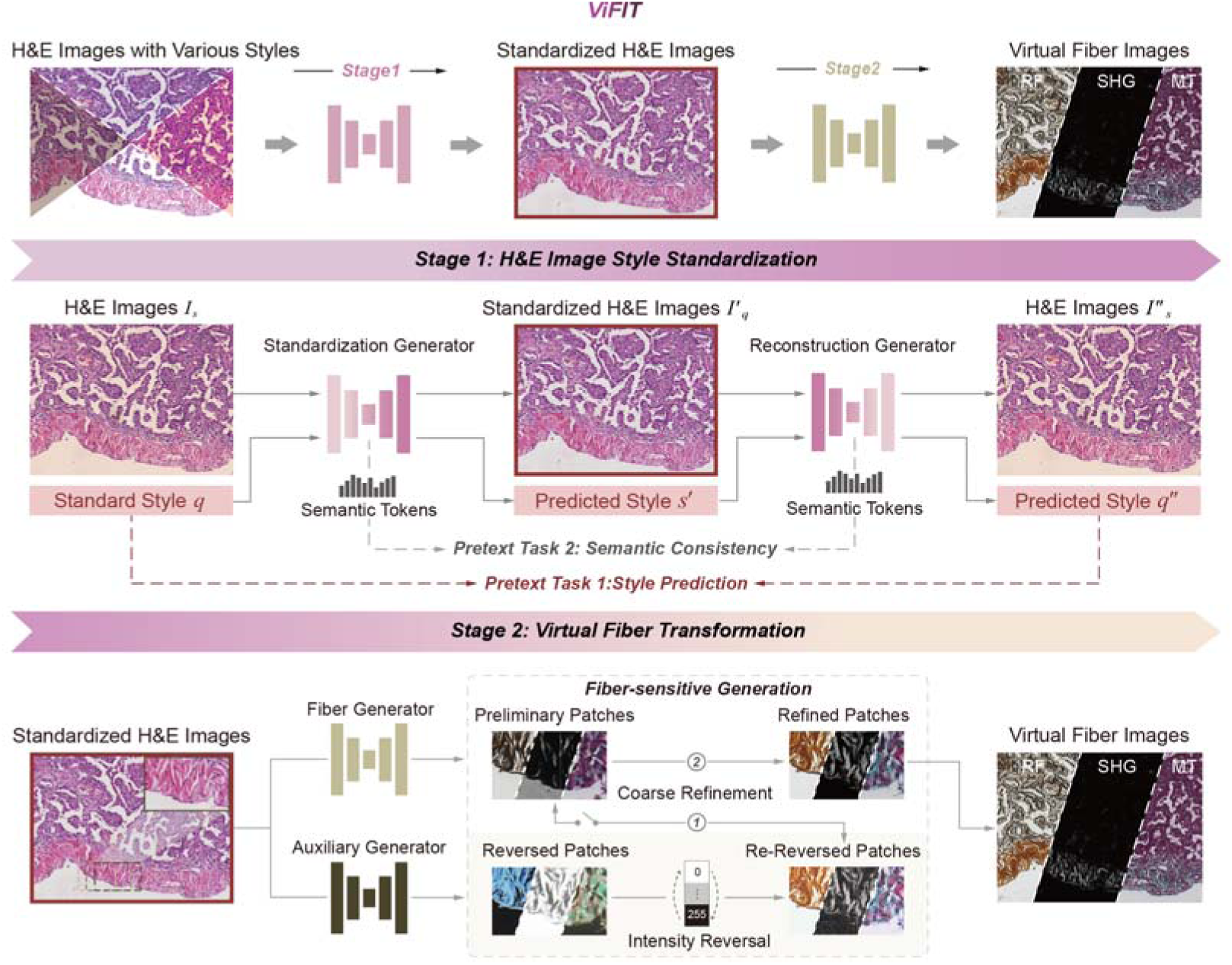
Workflow of the proposed ViFIT. ViFIT operates in two stages: H&E image style standardization (Stage 1) and virtual fiber transformation (Stage 2). In Stage 1, ViFIT employs dual generators—the Standardization Generator (S-Gen) and the Reconstruction Generator (R-Gen)—to transform H&E images of varying styles into a unified, standardized style. During the cyclic training process (one of the cascading orders for two generators is illustrated above), two pretext tasks are introduced for each generator: 1) incorporating the query style as generation guidance while predicting the input image’s style; and 2) extracting its semantic tokens for preserving the semantic consistency of generators’ features. In Stage 2, the standardized H&E images serve as the foundation for generating diverse fiber images. An unsupervised virtual fiber transformation network is employed, featuring a coarse-to-fine process. This process comprises of the generation of preliminary images through an intensity-reversed fiber-sensitive generation scheme and the refinement of coarse fiber images.

To effectively transfer the H&E image style to the standard one, the key is to disentangle the image content and styles within the extracted features. This process aims to enhance the representativeness of style features while ensuring that the image content remains unaltered during generation. Hence, we innovate the style generator by adding two pretext tasks, style label prediction and semantic consistency preservation. To achieve the first pretext task, as shown in Stage 1 of Fig. 1, each generator (including S-Gen and R-Gen) is provided with an extra input, i.e., the query style label that guides the generator to produce the H&E image with the particular style, and assigned with an additional output, specifically for predicting the style label of the input H&E image. This design leads to a tandem optimization scheme, as it introduces two effective constraints: 1) it requires the second generator to correctly predict the style of the images generated by the first generator; 2) it uses the style label predicted by the first generator as the query style label for the second generator, thereby guiding the style of its generated images. On one hand, if the first generator fails to generate images with the desired style, the second generator will not be able to correctly predict its style label. On the other hand, if the first generator fails to correctly predict the style label, it would severely affect the reconstruction quality of the second generator. In this way, the first pretext task effectively motivates both generators to learn the representation of image styles. As the second pretext task, we design a side network branch for each generator to extract semantic tokens that disentangle image content from the encoded features. As illustrated in Fig. 1, we ensure that the semantic tokens of the cascaded generators remain consistent, gradually extracting the similar underlying components of the encoded features that represent the image content, thereby preserving it unchanged during the generation process.

In the second stage, the goal is to transform a standardized H&E image into the desired fiber images under the guidance of unpaired H&E images and fiber images. A typical GAN-based model tends to introduce artifacts during image generation, due to the lack of ground truth for supervising the adversarial training process, resulting in low-quality and unreliable images. To overcome this limitation, we present a novel fiber-sensitive generation scheme that first utilizes image intensity reversal to draw the attention of the model to fiber structures in a self-supervision manner and thus progressively strengthens the quality of the generated fiber images (please refer to the Stage 2 of Fig. 1). To achieve this goal, in addition to a typical fiber generator, we introduce an auxiliary generator, dedicated to generate the intensity-reversed fiber image, which facilitates the model to concentrate on the fibrous tissue regions. To this end, the intensity-reversed patches generated by it can serve as pseudo ground truth. By re-reversing their intensity, we can supervise the fiber generator to produce preliminary patches that clearly delineate fiber foreground information. Once the fiber generator has learned to produce these preliminary patches from the auxiliary generator, the auxiliary generator is paused, allowing the fiber generator to refine the fiber image generation further.

### 2.2 H&E-stained Image Style Standardization

To validate the H&E style standardization performance of ViFIT, we used the images with five different representative H&E styles from our collected dataset and BreakHis public dataset^24^ as input images (Fig. 2). Fig. 2a illustrates the transformation of a large-scale breast cancer H&E image, exhibiting color deviation, into a standardized style image. The standard style was defined it as the consensus-driven style established by 16 pathologists from different institutions (detailed in a user study in Supplementary Note 1 and Table 1), which serves as the ground truth (GT) during the H&E staining standardization stage.

**Fig. 2.**
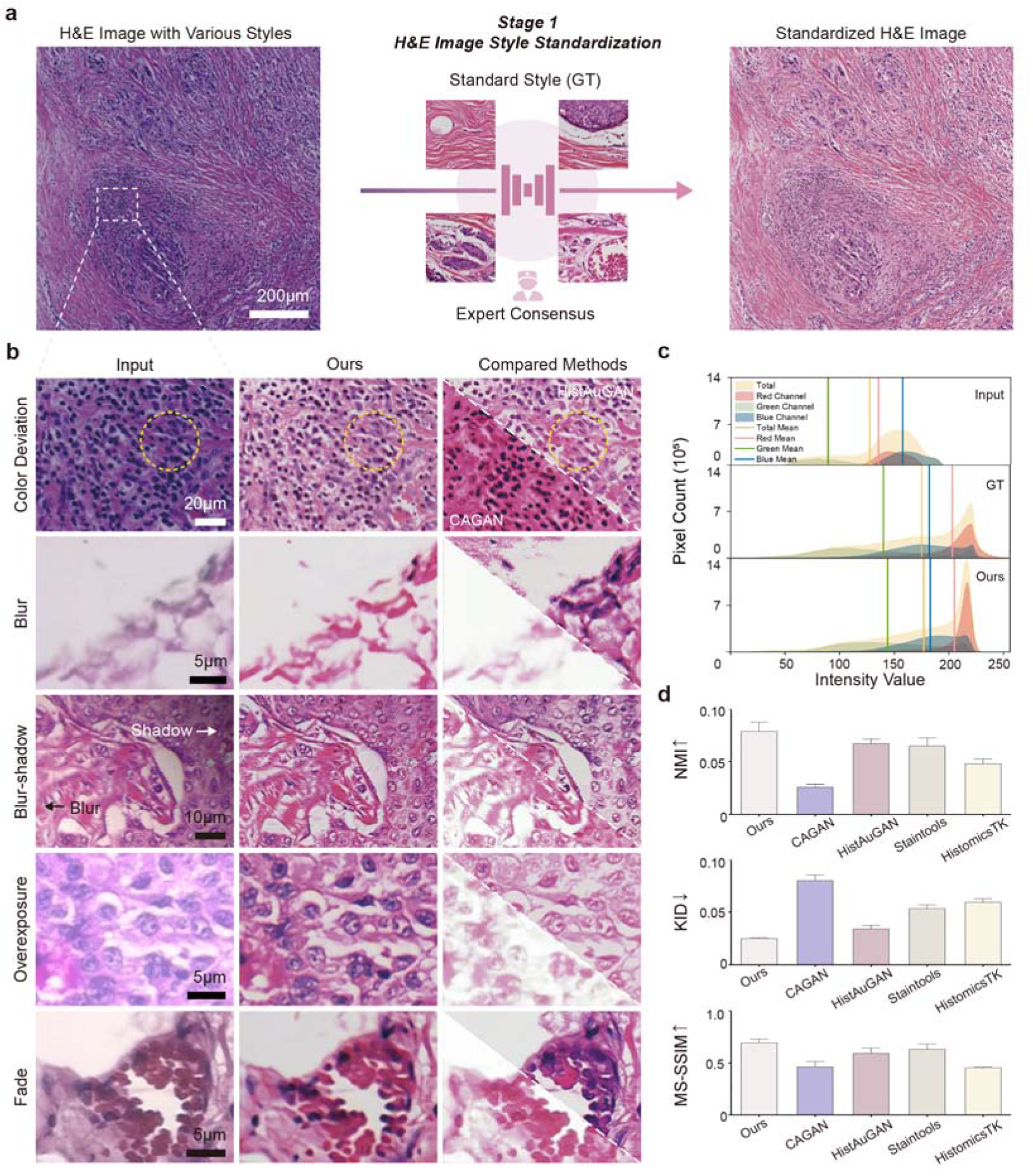
Style standardization of H&E-stained images with diverse style variations using ViFIT. **a** Illustration of the H&E image style standardization process for a large-scale breast cancer image exhibiting color variation. **b** Magnified standardization results of five typical H&E image styles, selected from our collected dataset and BreakHis public dataset. Yellow circles highlight regions where nuclei were mistransformed by the compared methods, while our method correctly performs the transformation. **c** Visualization of intensity distribution across RGB channels for the image in **a**, illustrating that ViFIT exhibits RGB distributions closely resembling the ground truth (GT). **d** Quantitative comparison demonstrates that, compared to off-the-shelf H&E style standardization methods, our method achieves higher NMI and MS-SSIM values (approaching 1) and lower KID values (nearing 0), indicating greater similarity of the standardized results to the GT (*n* = 162, where *n* represents the number of regions of interest (ROIs) evaluated for standardization).

In Fig. 2b, we zoom in on representative styles, including color deviation, blur, blur-shadow, over-exposure, and fading. These variations arise not only from slide quality but also from factors such as defocusing or detector noise during scanning in digital pathology systems^25,26^. Comparing to the other unsupervised style standardization models, CAGAN_27_ and HistAuGAN_28_, ViFIT effectively captures the color, contrast, and brightness of standard style, while accurately preserving both cellular and extracellular matrix features. By contrast, CAGAN exhibits substantial deviations from the standard style. Although HistAuGAN corrects color-deviation images to bring them closer to the GT, the yellow circles in Fig. 2b highlight its misclassification of nuclei, often falsely generating collagen fibers or blood cells instead, significantly reducing the number of identifiable nuclei. For the other styles, HistAuGAN also confuses nuclei with other tissue components, frequently misinterpreting the white background as collagen fibers. This error could be attributed to the varying tissue characteristics across different H&E staining styles. Notably, for the blur and blur-shadow images, ViFIT not only corrects the blurred regions (black arrow) but also restores the shadow areas (white arrow), demonstrating its unique image enhancement capabilities. Furthermore, we compared two conventional digital image processing-based H&E image standardization tools, Staintools^29^ and HistomicsTK (https://github.com/DigitalSlideArchive/HistomicsTK). These methods primarily focus on overall color correction, making it challenging to adapt to diverse stylistic variations (Extended Data Fig. 1). Additionally, in consideration of pathologists’ viewing preferences, we also demonstrate that ViFIT can transform diverse H&E images into other alternative staining styles suitable for diagnostics (Extended Data Fig. 2).

For evaluation, we first visualized the intensity distribution across RGB channels for the color-deviated H&E images (Fig. 2c). ViFIT demonstrates robust color preservation capabilities, with its color distribution and mean intensity values closely aligning with those of the GT, outperforming the other comparative methods (Extended Data Fig. 3). Next, we quantified standardization performance using normalized mutual information (NMI)^30^, kernel inception distance (KID)^31^, and multi-scale structural similarity index measure (MS-SSIM)^32^, which assess style clustering, tissue characteristics, and multi-scale structure. ViFIT outperforms four comparative methods across all three metrics. Although HistAuGAN exhibits some capabilities in standardization, its performance is approximately 10% lower than that of ViFIT. In addition, we conducted a user study to subjectively score the standardized results of the aforementioned methods (Supplementary Note 2 and Table 2). Both the quantitative evaluations and user study consistently demonstrate ViFIT’s superior performance in standardizing H&E staining styles.

### 2.3 Virtual Fiber Transformation Based on Standardized H&E

To evaluate the virtual fiber image transformation capabilities of ViFIT, we utilized the standardized H&E images of invasive lobular carcinoma and invasive ductal carcinoma of the breast as input, transforming them into SHG collagen and RF-stained images, respectively (Fig. 3). We compared our approach with state-of-the-art unsupervised virtual staining models, UTOM^20^ and VirtualMultiplexer^22^, as well as image transformation models, DRIT^33^ and UVCGAN^34^. For a fair comparison, all models were fed with the same standardized H&E images. Our results demonstrate superior SHG transformation performance in preserving histopathological features. In contrast, VirtualMultiplexer fails to generate a complete set of collagen fibers (arrows in Fig. 3a), while UTOM mistakenly generates cells as collagen fibers (circles in Fig. 3a). DRIT partially transforms collagen fiber but lacks sufficient intensity, whereas UVCGAN struggles to generate valid image content. Overall, ViFIT precisely generates collagen fiber tissue without extraneous branches, facilitating clearer interpretation by pathologists. Notably, SHG signals are predominantly generated by non-centrosymmetric collagen fibers^35,36^. Since the reference SHG images were obtained from adjacent unstained tissue sections, our results closely match the reference and clearly outperform competing methods. Furthermore, in Extended Data Fig. 4, we validated the reliability of collagen fiber generation results produced by ViFIT through a comparison with adjacent MT-stained sections.

**Fig. 3.**
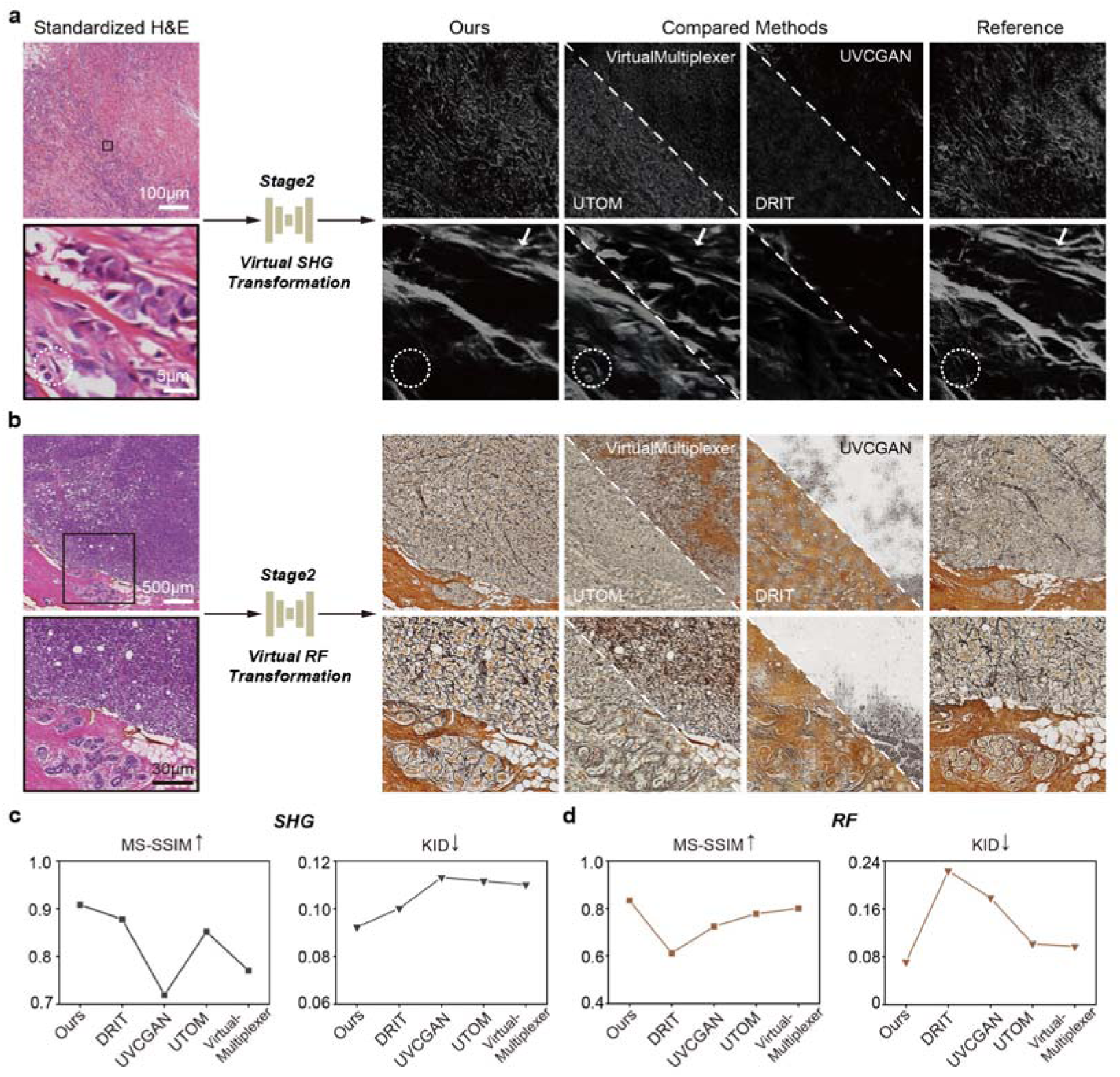
Virtual transformation of collagen and reticular fiber using ViFIT. **a** Virtual transformation of SHG collagen fiber images based on the standardized H&E images obtained from the previous stage. In SHG images, white structures correspond to the pink collagen fibers in H&E images. Compared to the state-of-the-art unsupervised virtual staining models, VirtualMultiplexer fails to synthesize continuous and complete collagen fibers (arrows), and UTOM produces the nuclei mistransformations (circles). **b** Virtual transformation of RF-stained images based on the standardized H&E images. ViFIT accurately captures both reticular and collagen fiber features, with yellow structures representing collagen fibers and brown ones indicating reticular fibers. **c** and **d** ViFIT outperform other methods, as demonstrated by the MS-SSIM and KID metrics, which quantify the similarity between the generated images and reference fiber images (*n* = 200, where *n* represents the number of ROIs evaluated for image transformation).

In the RF-stained images (Fig. 3b), reticular fibers are depicted in black, whereas collagen fibers are shown in brownish-yellow^37^. ViFIT accurately captures the features of both reticular and collagen fibers, while other models struggle to differentiate between these two fiber types, leading to significant clustering errors, such as misidentifying reticular fibers as collagen fibers. Quantitatively, we evaluated the similarity between the ViFIT-generated fiber images and their corresponding reference images using MS-SSIM and KID metrics (Fig. 3c and d). ViFIT outperforms all comparison methods, surpassing DRIT by 18% in virtual SHG evaluation and exceeding VirtualMultiplexer by 20% in virtual RF evaluation. These results demonstrate that, with style-standardized H&E images as input, ViFIT achieves superior performance in fiber image transformation. Furthermore, we conducted experiments to highlight the necessity of the H&E image style standardization stage, which significantly reduced the amount of training data required for generating virtual fibers (Supplementary Fig. 1).

### 2.4 ViFIT’s Adaptability Across Different Diseases

The capability of ViFIT arises from the integration of image style standardization and virtual fiber transformation. As shown in Fig. 4, we validated ViFIT’s performance across the entire workflow using H&E images from breast cancer, lung cancer, and colorectal cancer. Specifically, we present both the standardized H&E images produced in the first standardization stage and the fiber images generated in the second transformation stage. Compared to the reference images from adjacent stained sections, ViFIT produces reasonable fiber images despite tissue variations across different diseases. Without the “standardization-transformation” dual-stage process, the other unsupervised methods exhibit fiber-cell misclassification. In Extended Data Fig. 5, when unstandardized H&E images are input into the virtual fiber transformation network, misclassification of histopathological features is avoided, but the resulting weaker signal intensity suggests that some fibers and tissue structures may not be accurately generated. These findings highlight not only the positive impact of H&E standardization on the accuracy of subsequent virtual fiber transformation, but also demonstrate ViFIT’s adaptability across diverse diseases.

**Fig. 4.**
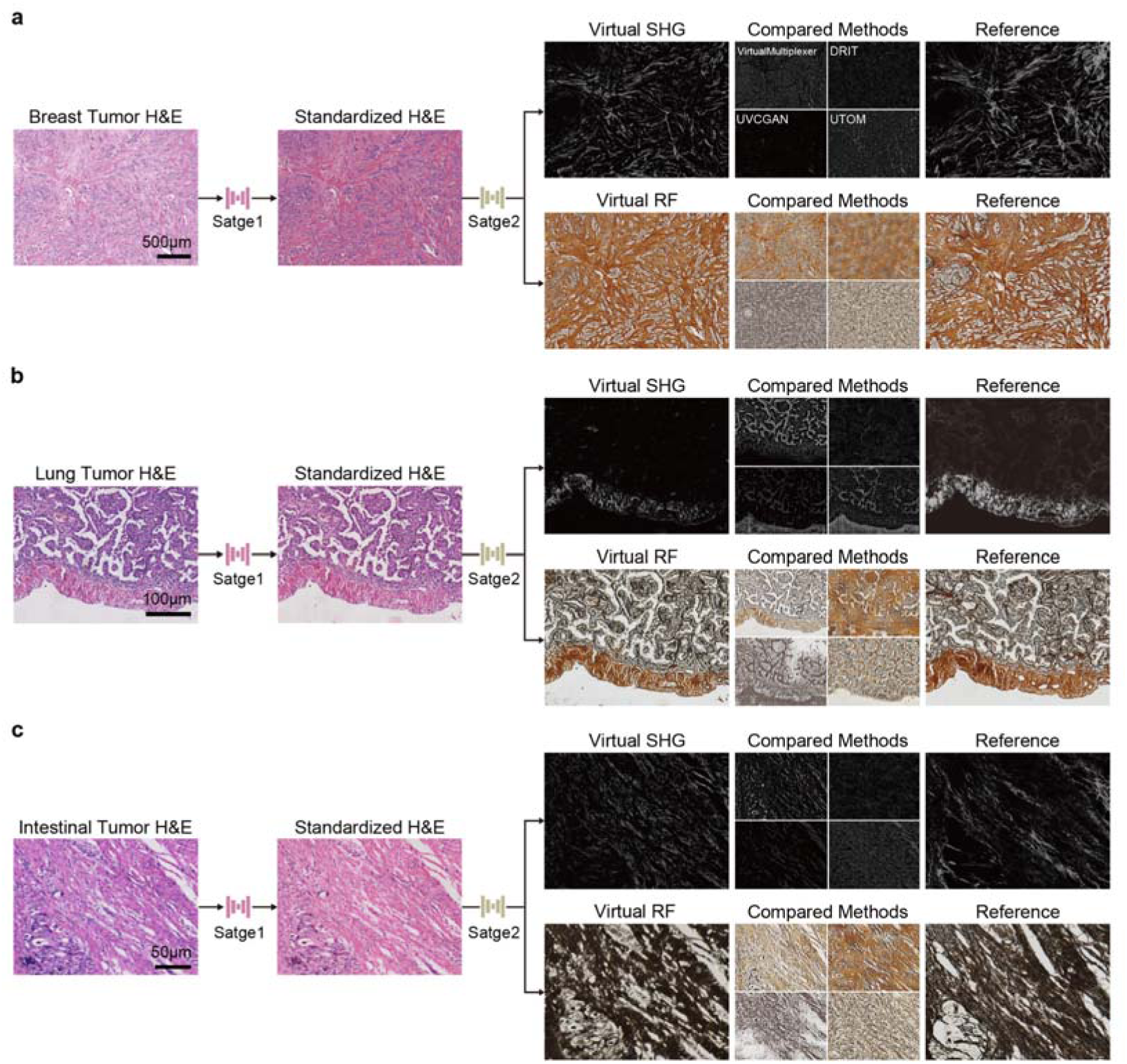
ViFIT generates fiber images across different diseases. **a**, **b** and **c** illustrate the adaptability of ViFIT to H&E images of breast, lung, and intestinal tumor, respectively. The “standardization-transformation” dual-stage process of ViFIT enables robust conversion of diverse H&E style images to the target SHG and RF-stained images. In contrast, comparative methods frequently struggle to properly produce the desired fiber images across different H&E staining styles.

To further validate the feasibility of the generated virtual fiber images for downstream quantitative analyses, we conducted fiber quantity, density, and alignment assessments using the CurveAlign software^38^, as shown in Fig. 5a. Additionally, perceptual similarity was evaluated through the VGG-16 network^39^, with representative breast SHG and RF images visualized in Fig. 5b and c. Notably, the fiber distribution and alignment in the virtual fibers generated by our method demonstrate highly consistent with the corresponding reference images. Both shallow texture features and deep perceptual analyses reveal that our model closely resembles the reference images, whereas virtual fibers generated by other models exhibit substantial discrepancies. For example, SHG images generated by UTOM and VirtualMultiplexer display an overabundance of well-aligned fibers with erratic collagen orientations, alongside positional inaccuracies in deeper perceptual regions.

**Fig. 5.**
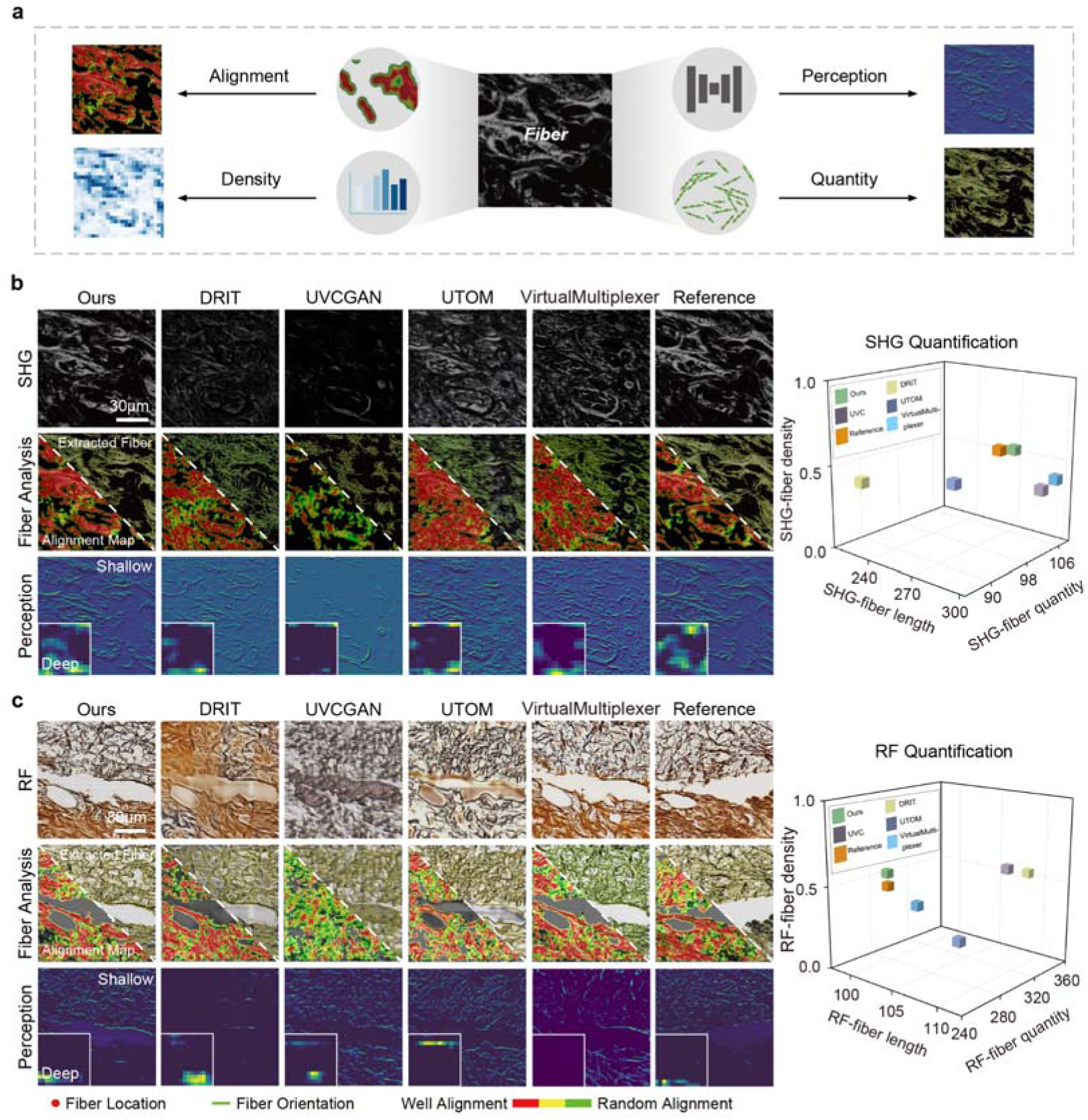
Downstream quantitative analyses of virtual fiber images. **a** The virtual fiber images generated by ViFIT are evaluated on the aspects of fiber quantity, density, alignment and perception similarity using CurveAlign software and the VGG-16 network, respectively. **b** and **c** visualize the fiber analysis and perceptual features of the breast cancer virtual SHG and RF images. The second row visualizes fiber location, orientation, and alignment, while the third row illustrates shallow and deep perception features. On the right of each row, the quantitative results, including fiber length, quantity, and intensity, highlight the accuracy of ViFIT in generating fiber images (*n* = 230, where *n* represents the number of ROIs evaluated for image transformation). Compared to other methods, the quantitative results of ViFIT-generated images most closely match those of the reference images across these downstream analysis tasks.

In addition, we visualized the quantification results of the fiber quantity, density, and length form SHG and RF images on the right side of Fig. 5b and 5c. As observed, for SHG quantification, the mean difference in fiber density between ViFIT and the reference images was limited to 0.03, while discrepancies in other models exceeded 0.1. Similarly, for RF quantification, ViFIT’s fiber length analysis aligns most closely with the reference image, both averaging 100, contrasting with deviations in other networks that reached up to 110. To sum up, these results underscore the robust accuracy of ViFIT’s fiber transformation, supporting its application in downstream tasks.

### 2.5 ViFIT-assisted Virtual Histopathological Diagnosis

The presence, rupture, and integrity of fibers are critical features indicative of hyperplasia or tumor^3,40^. To evaluate the potential of ViFIT in clinical pathological diagnostic workflow, we validated its diagnostic efficacy using five typical postoperative and intraoperative pathological cases (Fig. 6).

**Fig. 6.**
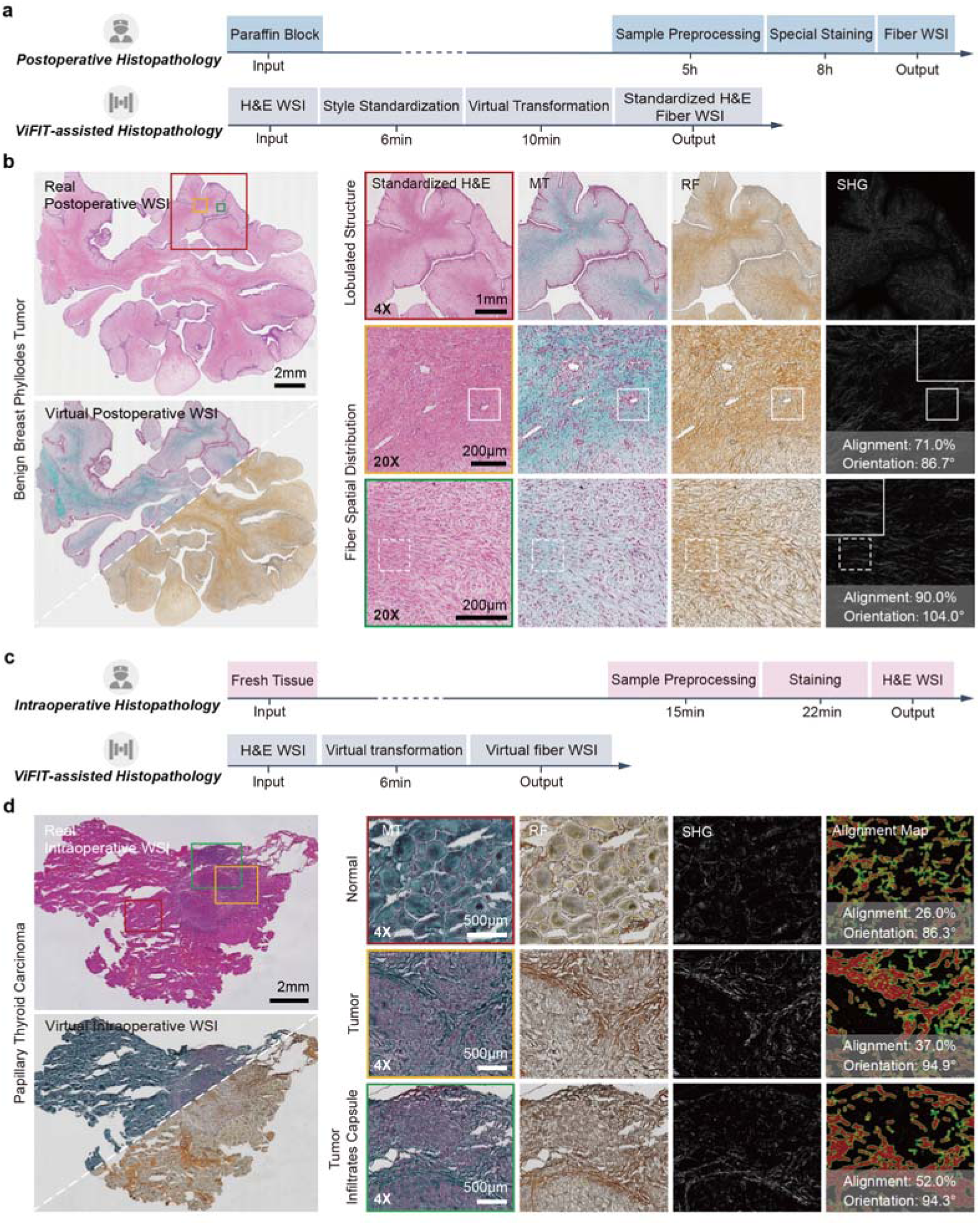
ViFIT-assisted histopathological diagnosis. **a** and **c** Comparison of postoperative, intraoperative, and ViFIT-assisted histopathology workflows. **b** ViFIT, based on standardized H&E images alongside virtual MT, RF, and quantitative SHG images, reveals mesenchymal-differentiated structures in postoperative breast tumor. **d** ViFIT enhances the accuracy of infiltration assessment in intraoperative thyroid carcinoma.

In the postoperative pathological workflow (Fig. 6a), obtaining special-stained fiber images typically requires over 5 hours of sample preprocessing by pathologists. In contrast, ViFIT, leveraging only digital H&E-stained images, generates both standardized H&E images and precisely registered virtual fiber images in just 10 minutes. As shown in Fig. 6b, ViFIT generates standardized H&E images alongside virtual MT, RF, and SHG images, capturing mesenchymal structures in benign breast phyllodes tumor at the whole-slide image (WSI) level, with high consistency to real-stained images (Extended Data Fig. 6a). At different magnifications, ViFIT accurately depicts fiber components within each lobule, including their spatial arrangement, alignment, and orientation relative to vascular structures (solid box) and cellular components (dotted box). For meningioma (Extended Data Fig. 7), ViFIT delineates tumor microenvironment features, emphasizing the relationship between fibers and native dural structures, while revealing heterogeneity in collagen and tumor distribution. In pituitary neuroendocrine tumors, ViFIT identifies collagen pseudo-capsules around tumor cell clusters, enhancing diagnostic precision for tumor boundaries (Supplementary Fig. 2).

In intraoperative frozen pathology, where time constraints often preclude special fiber staining, ViFIT achieves virtual fiber imaging within 6 minutes—well within the 25-minute diagnostic window. For instance, in papillary thyroid carcinoma (Fig. 6d), ViFIT effectively visualizes normal thyroid follicular structures (red box) and pronounced desmoplasia in the tumor stroma (yellow box). When tumor tissue infiltrates the capsule (green box), ViFIT enhances the accuracy of infiltration assessment by quantifying differences in collagen orientation and alignment. By contrast, benign basal cell adenoma of the parotid gland exhibits neither significant capsule invasion nor desmoplastic response (Extended Data Fig. 8).

Additionally, ViFIT-assisted histopathology significantly mitigates nonspecific staining artifacts caused by colloid deposition, slide wrinkling, tissue loss, or blotting (arrows in Extended Data Fig. 6b). It also preserves collagen network integrity under tissue compression, ensuring reliable visualization (Supplementary Fig. 2). These capabilities not only enhance staining quality but also enable low-magnification tissue architecture analysis and high-magnification fiber characterization. Overall, ViFIT redefines fiber staining workflows by offering cost-effective precision, rapid processing, and artifact suppression. Its capabilities, limitations, and time efficiency are summarized in Supplementary Tables 3 and 4, highlighting its transformative potential in clinical pathology.

## 3. Discussion

Special staining techniques and label-free SHG imaging provide complementary approaches for characterizing fibers from different perspectives^6,41^. . However, their diagnostic utility is limited by practical challenges, including the complex protocols required to obtain fiber-stained images from a single tissue slide. Additionally, the high cost and limited clinical adoption of SHG imaging systems hinder their widespread implementation in routine pathology. To overcome these barriers, we introduce ViFIT-assisted histopathology, which generates diverse fiber images and standardized H&E-stained images, facilitating seamless integration into diagnostic workflows.

The quality of H&E slides is paramount for accurate diagnosis. However, variations in staining reagents, protocols, and slide preparation across institutions introduce stylistic differences, not only between hospitals but even within the same hospital across different batches. This variability often necessitates resectioning and restaining of original paraffin-embedded tissues when patients present non-standardized slides at higher-level hospitals. Such practices increase diagnostic workloads and delay treatment decisions.

Our proposed unsupervised dual-stage framework addresses this issue by producing style-standardized H&E images as an intermediate output. Unlike conventional GAN-based models, our style standardization model employs two pretext tasks: style label prediction and semantic consistency preservation. Together, these tasks collectively enhance style representation and maintain image content integrity. For style label prediction, a tandem optimization scheme is introduced to the cyclic cascading structure of S-Gen and R-Gen, where S-Gen identifies diverse input styles, and R-Gen equipes with knowledge of the standard style. Misaligned predictions by either generator may disrupt the cyclic process, emphasizing the importance of precise style identification. Thus, this approach prioritizes style prediction, thereby enhancing the generators’ ability to represent styles accurately. For semantic consistency preservation, we aim to address the challenge of style-content interference, as style transformations often introduce artifacts or misclassify tissue types. The primary reason is that the style and content components are entangled within the encoded features, making style transformation likely to alter the histopathological features. To mitigate this, we extract semantic tokens from encoded features using side network branches. By extracting and enforcing consistency of semantic tokens during training, our model separates style from content, preserving histopathological features and ensuring artifact-free transformation.

In essence, the variability in H&E staining styles introduces fine-grained complexity in the representation space, posing significant challenges for existing end-to-end frameworks that convert diverse H&E images into specific fiber images. These frameworks rely heavily on extensive paired datasets and intricate models to accommodate heterogeneous data distributions, complicating the development of robust image generation models. In contrast, our unsupervised ViFIT framework mitigates this challenge by standardizing input styles, reducing the dimensionality of the representation space and simplifying the training process for virtual image transformation.

Therefore, ViFIT-assisted histopathology offers several distinct advantages. First, it addresses the scarcity of paired H&E-to-fiber image datasets. While large datasets of H&E images are readily available for style standardization, acquiring equivalent volumes of fiber images is significantly more challenging. By employing a two-stage unsupervised design, ViFIT enables robust standardization using widely accessible H&E image datasets, followed by fiber image transformation with a relatively small H&E-to-fiber dataset (Supplementary Fig 1). Second, ViFIT facilitates simultaneous cross-modal generation of label-free and stained fiber images via a novel intensity-reversal strategy. Differences in fiber color and intensity between modalities complicate the accurate translation of fiber locations from H&E-stained images to other fiber images. For instance, H&E fibers appear as high-intensity pink pixels, while SHG fibers appear as low-intensity white pixels. Directly mapping these modalities increases task complexity. Inspired by curriculum learning^42^, we employ an auxiliary generator to produce intensity-reversed patches, allowing the model to focus on fiber structures during early learning stages and subsequently refine details in later stages, thereby improving the accuracy of fiber transformation.

Validated across diverse intraoperative and postoperative cases, ViFIT streamlines slide review processes and complements established clinical pathological workflows. Despite these benefits, the limitations of our proposed approach are as follows. Currently, generating each type of fiber image requires training an independent model, which increases computational burden. Future research could explore the possibility of making this model more versatile, capable of producing multiple fiber images within a single model. As larger H&E-to-fiber datasets become available, a fiber transformation foundation model could enable incremental adaptability to diverse H&E styles. Furthermore, incorporating text prompts for interactive customization could provide tailored outputs, enhancing diagnostic flexibility.

## 4. Methods

### 4.1 Ethical Statement

All anonymous tissue collections for retrospective study were conducted under a protocol approved by the Institutional Review Boards of the First Affiliated Hospital of Fujian Medical University (2023-289) and Fujian Medical University Union Hospital (2020-085). For the user study, all the participants have provided written the informed consent prior to participating in the image style questionnaire.

### 4.2 Microscopy Datasets

We established two public unpaired image datasets: a multi-style breast H&E-stained dataset and an H&E-to-fiber dataset. The multi-style breast H&E-stained dataset comprises 8,578 H&E-stained images representing seven distinct staining styles, with varied sizes and magnifications. These images were sourced from our collection at Fujian Medical University Union Hospital, along with publicly available datasets, including BreakHis^24^, BACH^43^, and BreCaHAD^44^.

The H&E-to-fiber dataset consists of H&E-stained images alongside corresponding RF, MT, and SHG images obtained from adjacent tissue slices. Notably, the SHG images were acquired using a multiphoton imaging system^45^ on unstained sections, with an excitation wavelength of 810 nm and an average power of 30 mW. This dataset includes 112 images of diverse sizes and magnifications, including WSIs, sourced from The First Affiliated Hospital of Fujian Medical University and Fujian Medical University Union Hospital. The dataset covers seven distinct tissue disease types: breast tumor, lung tumor, intestinal tumor, meningioma, pituitary neuroendocrine tumor, thyroid tumor, and basal cell adenoma of the parotid gland. Further details regarding these datasets are provided in Supplementary Tables 5 and 6, which summarize the source institutions, specimen types, imaging modalities, image counts, resolutions, and magnification levels. The datasets will be publicly released after accepted.

### 4.3 H&E-stained Image Style Standardization

To achieve the transformation from H&E images to virtual fiber images, we formulate it as a standardization-transformation dual stage process. In the H&E image style standardization stage, a cyclic generative adversarial network is introduced to facilitate fundamental style conversion from different staining styles to a standard style. As shown in Supplementary Fig. 3, the proposed H&E image style standardization model achieves the purpose through the following components: (1) two ResNet-based generators (S-Gen and R-Gen), which are trained cyclically to perform style standardization; (2) CNN-based classifiers as discriminators, utilized to regionally differentiate the original and query style images; (3) a cycle consistency loss, which ensures that original or query style images generated by the generator can be restored to their original form. On this basis, we introduce two effective pretext tasks to facilitate unsupervised learning.

#### Preliminaries

We set up a pool of style labels, denoted as **S** = {0,…,*S*}, based on their respective sources. Note that the label of the standard style is set as 0. For a standard generator, given the input image *I*, it will generate a new image with a different style, *I*’. The process can be briefly described as *I’* = *G*(*I*), which is supervised by an adversarial loss.

#### Pretext task 1: style prediction

In contrast to the standard generators, our proposed generators are assigned with a pretext task, style prediction. Specifically, each generator has dual inputs: an H&E image with a certain style *s*, *l_s_(l_s_* ∈ *R^H^*^×*W*×3^)and a query H&E image style *q (q* ∈ **S**) that guides the style of the generated image. It has two corresponding outcomes, a generated image with the query style, denoted as 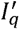, and the predicted style label of the input image, denoted as *s*’. Thus, the computation of a single generator *G* can be expressed as: 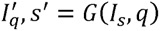.

On the training phase, two generators with identical structure are cascaded and assigned with different roles. S-Gen, denoted as *G_S_,* is dedicated to transform the H&E image to the standard style. R-Gen, denoted as *G_R_,* aims to recover the standard style image to an image with the given style. Generally, the cyclic computing process of the cascaded generators, including the cascading orders from S-Gen to R-Gen *(i.e.,S → R)* and from R-Gen to S-Gen *(i. e.,R → S),* can be described as below, respectively.

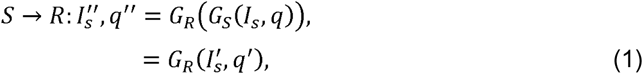

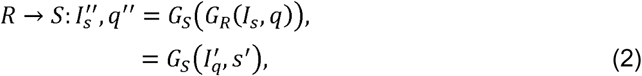

where 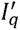 and *s’* represent the outcomes of the first generator. 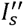 and *q”* refer to the results of the following generator. For instance, for *G_S_* in S → *R*, *l_s_* is a sampled H&E image with a particular style *s* and *q* represents the standard style, i.e. 0, as query. In this way, we utilize the cyclic training framework and the pretext task to develop a tandem optimization scheme, as shown in Fig. 1. Under this scheme, not only the generated image from the first generator is used for the input of the second generator, but also the predicted style label from the first generator is utilized as the input query style for the second generator.

In practice, for the sake of computational efficiency, we expand into a matrix with the same dimension as that of *l_s_* and append it to *I*_s_, i.e., *T =* {*l_s_,**Mat***(*q*)}(*T* ∈ *R^H^*^×*W*×4^) which is fed into the generator as input, so that the generator can output a tensor *T” (T”* ∈ *R^H^*^×*W*×4^) where the first three channels correspond to 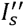 s and the last channel correspond to the style label prediction. To this end, the training objective can be described as below.

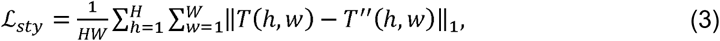

where *T(h,w)* and *T”(h,w)* represent the vectors at the position (*h,w*) (*h* ∈ {1*,…,H*}), *w* ∈ {1*,…,W*} of the tensor *T* and *T”*.

#### Pretext task 2: semantics consistency

We introduce an additional pretext task to preserve the image content unchanged during generation. To achieve this, we derive what we call ‘semantic tokens’ from the encoded features of each generator. The tokens are obtained through multiple convolutional layers and a multi-perceptual layer, which compactly describe the content of the input H&E images. Subsequently, we encourage the semantic tokens of the generators to be consistent as the objective of the pretext task. Specifically, the tokens of S-Gen and R-Gen are denoted as *x_S_* an *x_R_,* so the objective can be defined as:

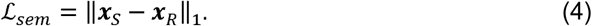

In overall, the training objective of the proposed style standardization model can be expressed as:

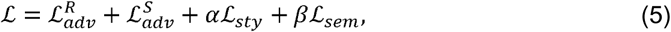

where 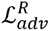 and 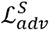 refer to the adversarial losses of S-Gen and R-Gen, which require unpaired images for style supervision. α and β represent the balancing weights for the pretext tasks, which is set as 60 and 12 in practice.

### 4.4 Virtual Fiber Transformation

In the virtual fiber transformation stage of ViFIT, the goal is to transform a standardized H&E image into the desired fiber images under the guidance of unpaired exemplar fiber images. As shown in Supplementary Fig. 4, we employ two generators: the fiber generator and the auxiliary generator, to create various collagen fiber images. Specifically, the fiber generator is primarily responsible for generating virtual fiber images from the standardized H&E images, while the auxiliary generator supplements the images produced by the fiber generator with salient fiber information. In practice, both generators are based on ResNet^46^ architecture. Additionally, CNN-based discriminators and cycle consistency loss are also utilized to provide adversarial supervision.

#### Intensity-reversal fiber-sensitive generation

The major challenge of generating fiber images lies in the indistinguishability of fiber structures, cellular components, and backgrounds. To address this issue, we discovered that reversing the pixel-wise intensity of fiber images provides a surprisingly simple yet effective way to visually separate fibers from the background. In the light of this, we introduce an intensity-reversal fiber-sensitive generation scheme, in which the auxiliary generator is tasked with generating intensity-reversed images rather than the normal fiber images, enabling the model to focus on the fiber regions. Formally, given the input image *I*_H&E_, the auxiliary generator *G*_aux_ aims to produce the intensity-reversed fiber image 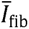 i.e., 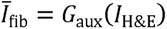, which is supervised by an intensity-reversed real fiber image for reference. Thus, the training objective of the auxiliary generator can be expressed as: 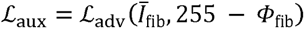, where 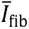 denotes the intensity-reversed image generated by the auxiliary generator. *G*_aux_ and *Φ*_fib_ represents the reference fiber image which is used to supervise the generation yet unpaired with the input H&E image. Subsequently, using the output from the auxiliary generator for coarse and preliminary supervision, we can correct the misinformation generated by the fiber generator and accurately preserve the fiber structures. To accomplish this, we flip the pixel values of the generated intensity-reversed images back to their normal chromatic domain, i.e., 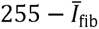 as pseudo ground-truth of the fiber generator.

#### Coarse refinement training strategy

Generally, the virtual fiber transformation model is trained in an unsupervised manner using unpaired H&E and fiber images. To enhance the fidelity of the generated images, we introduce a coarse refinement training strategy aimed at producing high-quality virtual fiber images. This approach facilitates a two-step training process. Initially, the fiber generator is optimized using adversarial loss and concurrently supervised by the re-reversed patches generated by the auxiliary generator. Consequently, the fiber generator gains the ability of producing preliminary patches that accurately preserve the fiber structures. Upon completing an epoch, we cease training the auxiliary generator and continue training the fiber generator, focusing on refining the details within the fiber patches. Thus, the training objective of the fiber generator *G*_fib_ consists of two adversarial losses:

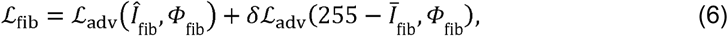

where 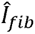 refers to the generated image of the fiber generator *G*_fib_ and 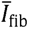 denotes the image generated by the auxiliary generator *G*_aux_. *Φ*_fib_ represents the reference fiber image. δ (δ ∈ {0,1}) is set as 1 at the initial stage of training, and set as 0 after training for an epoch. Furthermore, we performed ablation studies to demonstrate the effects of different components of ViFIT (Supplementary Note 3 and Supplementary Fig. 5).

### 4.5 Implementation Details

All generators, including those for H&E style standardization, the fiber generator, and the auxiliary generator, utilize the identical network architecture based on ResNet^46^ consisting nine residual blocks. Specifically, the style standardization model processes an H&E image of dimensions 256×256×3, along with a query style label that is expanded into a 2D matrix of size 256×256 and appended to the input image, resulting in an input tensor of size 256×256×4. Similarly, the model’s output comprises the combination of the H&E image with the desired style and the predicted style matrix, resulting in a tensor of dimensions 256×256×4. In the meantime, we apply two convolution layers with the kernel size 3×3×512 and a fully connected layer to compute a semantic token with the dimension of 1×10, which represents the style profile of the input H&E image. Additionally, the virtual fiber transformation model processes the standardized H&E images with the same dimension of 256×256×3 and generates a fiber image with the same dimension.

During the training phase, the Adam optimizer is used for all the networks. The learning rate is set to 0.0002, with the exponential decay rates for the first moment estimate and the second moment estimate set to 0.5 and 0.999, respectively. The total number of training epochs is 65, with a fixed learning rate for the first 5 epochs and a linear decay of the learning rate over the subsequent 60 epochs.

### 4.6 Comparison Methods

For evaluation, we validated the capabilities of ViFIT in both H&E-stained image style standardization and virtual fiber transformation stages, respectively. For style standardization comparison, we introduced two advanced open-source unsupervised models for H&E image style standardization, HistAuGAN^28^ and CAGAN^27^, as well as two conventional digital image processing-based H&E style standardization tools, Staintools^29^ and HistomicsTK. Specifically, HistAuGAN relies on a domain-content-preserving encoder to control style standardization, while CAGAN proposes a dual-decoder generator and force consistency constraints to standardize the style of H&E images. Staintools standardizes H&E images applies sparse non-negative matrix factorization to separate and normalize stain components. HistomicsTK standardizes H&E images through a Python package for the analysis of digital pathology images.

For virtual fiber transformation, we employed the latest unsupervised virtual staining model for microscope images, VirtualMultiplexer^23^ and UTOM^20^. Specifically, VirtualMultiplexer achieves virtual immunohistochemical staining with a multiscale constrained architecture at the single-cell, cell-neighborhood, and whole-image level. UTOM employs a content-preserving transformation module to ensure content coherence. Notably, these two unsupervised models require minimal expert intervention. To ensure a fairer comparison, we modified the local cell consistency loss in VirtualMultiplexer to eliminate reliance on expert cell annotations. For UTOM, we manually adjusted content thresholds to maintain the similarity between H&E domain and fiber domain.

Additionally, we compared two unsupervised style transfer models for natural images were presented for comparison, DRIT^33^ and UVCGAN^34^. DRIT uses a content discriminator to facilitate the factorization of domain-invariant content space and domain-spec45ific attribute space, achieving seasonal changes in natural landscapes. UVCGAN combines UNet and Vision Transformer, using UNet for high-frequency feature processing and Vision Transformer for low-frequency feature handling, enabling style transfer in facial transformations.

### 4.7 Evaluation Protocols

To evaluate the quality of ViFIT-generated images, we first define the GT or reference image tailored to each stage of the process. For the H&E image style standardization stage, the GT is a standardized style established through consensus among pathologists. In the virtual fiber transformation stage, the reference image is represented by real fiber images obtained from adjacent tissue slices.

We then apply several quantitative metrics—NMI^30^, KID^31^, and MS-SSIM^32^—to rigorously assess the generated images against these reference standards. Specifically, NMI measures style clustering accuracy, reflecting effectiveness in style standardization; KID quantifies the feature space distance between the generated images and the GT, utilizing kernel functions to assess fidelity without supervision; and MS-SSIM measures brightness, contrast, and structural similarity across multiple scales, providing a perceptual comparison. Additionally, we visualize the intensity distribution across the RGB channels to further evaluate the accuracy of the style standardization.

To further evaluate the performance of ViFIT in downstream fiber quantification, CurveAlign is employed to analyze fiber orientation, length, and quantity, generating images with extracted fibers and corresponding fiber alignment heatmaps. Additionally, the pre-trained VGG-16 network is utilized to extract both shallow texture features and deep perceptual features from the fiber images, enabling a comprehensive evaluation of fiber characteristics.

### 4.8 Hardware and Software

All experiments were conducted on a workstation with a single NVIDIA GeForce RTX 3090 GPU. Each model was implemented and trained using Python (v3.7.1) and PyTorch (v1.8.0).

## Data Availability

Our collected multi-style H&E-stained image dataset as well as the H&E-to-fiber dataset will be available upon acceptance. The BreakHis dataset are available at https://www.kaggle.com/datasets/ambarish/breakhis. The BACH dataset are available at https://iciar2018-challenge.grand-challenge.org. The BreCaHAD dataset are available at https://bmcresnotes.biomedcentral.com/articles/10.1186/s13104-019-4121-7.

## Code Availability

The codes, pretrained models, and relevant resources of the proposed ViFIT will be publicly released upon acceptance.

## Acknowledgements

We would like to thank Xi Chen and Jianping Huang for their support in the data acquisition, as well as Houqiang Li, Zongxian Ye, and Dehua Zeng for their assistance in defining the staining styles. This work was support by the National Natural Science Foundation of China (62005049 to S.W., 62072110 to W.L., and 82171991 to J.C.), the Natural Science Foundation of Fujian Province (2024J02013 to J.C. and 2024J01624 to D.K.), the Scientific Research Project of National Key Clinical Specialty Construction Project (2022YBL-ZD-03 to X.W.).

## Author Contributions

S.W., W.L., F.H., and J.C. designed and directed the study; C.L., S.W., and W.L. set up the ViFIT framework; X.Z., C.L., and X.L. performed quantitative analysis; S.W., and X.H. performed multiphoton imaging experiments; X.W., X.H., D.K., R.L., and L.H. provided specimen; S.W., W.L., X.Z., X.W., and C.L. wrote the paper with contributions from all the authors.

